# Novel bacteriophages targeting wheat phyllosphere bacteria carry DNA modifications and single-strand breaks

**DOI:** 10.1101/2024.10.22.619576

**Authors:** Peter Erdmann Dougherty, Maja Schmidt Pedersen, Laura Milena Forero-Junco, Alexander Byth Carstens, Jos M. Raaijmakers, Leise Riber, Lars Hestbjerg Hansen

## Abstract

The phyllosphere microbiome can positively or negatively impact plant health and growth, but we currently lack the tools to control microbiome composition. Contributing to a growing collection of bacteriophages (phages) targeting bacteria living in the wheat phyllosphere, we here isolate and sequence eight novel phages targeting common phyllosphere *Erwinia* and *Pseudomonas* strains, including two jumbo phages. We characterize genomic, phylogenetic, and morphological traits from these phages and argue for establishing four novel viral genera. We also search the genomes for anti-defense systems and investigate DNA modifications using Nanopore sequencing. In Pseudomonas phage Rembedalsseter we find evidence of 13 motif-associated single-stranded DNA breaks. A bioinformatics search revealed that 60 related *Pseudomonas* phages are enriched in the same motif, suggesting these single-stranded nicks may be widely distributed in this family of phages. Finally, we also search the Sequence Read Archive for similar phages in public metagenomes. We find close hits to the *Erwinia* jumbo-phage Kaldavass in a wide variety of plant, food, and wastewater metagenomes including a near-perfect hit from a Spanish spinach sample, illustrating how interconnected geographically distant phages can be.

## 2. Introduction

Although Earth’s surface is mostly ocean, the land surface area is dominated by plants, with vegetation covering 69% of the land surface area^1^. This extensive aerial vegetation is in turn colonized by bacteria, fungi, viruses, and other microorganisms that collectively constitute the phyllosphere microbiome^2^. The host plant often benefits from this microbial community through growth promotion^3^ or enhanced stress tolerance^4^, but the phyllosphere is also a major niche for plant pathogens^5^. For this reason, techniques capable of manipulating the phyllosphere microbiome are of great interest.

One such potential tool are bacteriophages (phages); viruses that infect bacteria^6^. Phages have attracted significant interest as tools for microbiome manipulation, either for the direct elimination of target bacteria (phage therapy/biocontrol)^7^, or through more indirect, broad-scale microbiome shifts (virome transplants)^8^. Broadly speaking, phage lifestyles can be split into two main types; virulent phages, which replicate immediately upon entering a viable host cell and kill the host soon after, and temperate phages, which can enter a dormancy state (prophage) within the host cell, retaining the ability to undergo induction and kill the host later^9^. Virulent phages are usually preferred for microbiome manipulation as temperate phages may integrate into new hosts, spreading phage resistance and possible virulence factors^7^.

To defend against bacteriophages, bacteria have evolved a tremendously diverse array of defense systems^10^ and in response, phages have evolved counter-defense systems^11^. One avenue of counter-defense is DNA modification, where phages add chemical groups to the canonical nucleotides and thus avoid recognition by some defense systems^12^. Although most modifications are undetectable with common DNA sequencing platforms (such as Illumina), their presence may be detected by sequencing non-amplified DNA on platforms such as PacBio or Oxford Nanopore Technologies (ONT)^13^. Recently, this approach was applied to human gut viromes, where an astonishing 97.6% of phages exhibited methylation^14^. However, more complex modifications may not be detectable in metagenomes, and screening for DNA modifications is not commonly performed for phage isolates (or viromes). We therefore know little about the distribution and frequency of phage DNA modifications in different environments.

In the phyllosphere, phages are highly diverse^15,16^ and have been shown to significantly alter bacterial abundances^17^. Recently, we also showed that some wheat phyllosphere bacteria harbor spontaneously induced prophages capable of killing rival bacterial strains, thus facilitating intra-species warfare^15^.

Although using prophage-infected bacteria to manipulate the phyllosphere microbiome is an intriguing idea, it does risk unwanted horizontal gene transfer. Approaching the problem from another angle, we have therefore previously isolated and characterized virulent phages against *Sphingomonas* strains from the same wheat phyllosphere as a first step in establishing a collection of phages targeting wheat phyllosphere bacteria^18^. Here, we continue this effort, isolating eight phages against wheat phyllosphere bacteria from two common phyllosphere genera, *Erwinia* and *Pseudomonas.* Sequencing revealed these phages to be highly diverse, with phylogenetic analysis supporting the establishment of four novel phage genera. We also conducted a bioinformatic search for anti-defense systems and investigated DNA structure and modifications in the phage genomes using ONT sequencing, identifying two methylated phage genomes, two phages with probable hyper-modifications, and one phage with single-stranded coverage indicative of motif-associated single-stranded DNA breaks (nicks). We find that a conserved group of *Pseudomonas* phages is enriched in the same nick-associated motif, and we investigate the distribution of this motif throughout their genomes.

## 3. Materials and Methods

### 3.1 Phage isolation and enrichment

The phages were isolated from household organic waste from a waste-management company (HCS A/S, Glostrup, Denmark). We have previously found organic waste to be a rich source for phages targeting plant-associated bacteria^18–21^. Following this established protocol, we centrifuged the organic waste samples at 5,000x*g* for 10 minutes followed by filtering with a 0.45 µm PVDF syringe filter (Merck Millipore, Germany). Seven *Erwinia* and *Pseudomonas* strains previously isolated from the wheat phyllosphere were used as bacterial hosts^15^. 100 µL of organic waste supernatant was added to 100 µL of the overnight bacterial cultures and plated with a double agar overlay with 4 mL LB soft agar (0.45% agarose, supplemented with 10 mM CaCl2 and MgCl2) on top of LB agar. After overnight incubation at 28 °C, plaques were picked, resuspended in SM buffer (100 mM NaCl, 10 mM MgSO4, 50 mM Tris-HCl, pH 7.5), and filtered using 0.22 µm syringe filters. To purify each phage isolate, plaques were picked and replated three times for each phage isolate.

High-titre (>10^8^ pfu/mL) phage enrichments were made by adding low titres of purified phage stock to exponential cultures of respective bacterial hosts in liquid LB media. The cultures were incubated overnight with shaking (225 rpm) at 28 °C, followed by centrifugation at 8000x*g* for 5 minutes to pellet the bacteria. The supernatants were then filter-sterilized as before.

### 3.2 TEM microscopy

To prepare phage samples for transmission electron microscopy (TEM) imaging, high-titre phage enrichments were purified by cesium-chloride (CsCl) gradient ultracentrifugation using an Optima XE-90 ultracentrifuge (Beckman Coulter, USA) with three CsCl density layers (1.3, 1.45, and 1.7 g/cm^3^). At the Electron Microscopy Facility at the University of Leicester, 5 µL of purified phage samples were spotted to a freshly glow-discharged carbon film grid for 2 minutes, washed with deionized water, and negatively stained with 1% uranyl acetate. Samples were then viewed on a JEM-1400 TEM (JEOL, Japan) with an accelerating voltage of 120 kV, with images taken using a Xarosa digital camera RADIUS software (EMSIS, Germany). Length and width values are the average of five measurements, shown with standard deviation.

### 3.3 DNA isolation and sequencing

DNA isolation was performed as in prior work^15^. Briefly, starting with high-titre phage enrichments, DNase I (25 units/mL) and RNase A (25 µg/mL) (A&A Biotechnology, Poland) were added to 500 µL of each phage amplification, and incubated for an hour at 37 °C. The nucleases were deactivated and encapsulated phage DNA released by incubation with EDTA (50µM), SDS (0.1 %), and Proteinase K (1 mg/mL) at 55 °C for an hour. Proteinase K was deactivated by incubation at 70 °C for 10 minutes. The raw DNA extractions were then purified with the Clean and Concentrator-5 kit (Zymo Research, USA). A Qubit 2.0 fluorometer with a High Sensitivity dsDNA assay (ThermoFisher, USA) was used to quantify the DNA concentrations. Illumina sequencing libraries were built using the NEBNext Ultra II FS DNA Library Prep kit (New England Biolabs, USA) according to the manufacturer’s protocol. Libraries were sequenced on a 151-bp paired-end sequencing run with the Illumina iSeq platform (Illumina, San Diego, USA).

To detect potential DNA modifications, the phage DNA was sequenced using the Oxford Nanopore Technologies (ONT) platform as previously described^22^. Briefly, a whole genome amplification (WGA) of each phage DNA isolation was prepared according to protocol using the illustra Ready-To-Go GenomePhi V3 DNA amplification kit (GE Healthcare, USA), followed by debranching with S1 nuclease (Thermofisher Scientific, USA). The product was then purified using the DNA Clean and Concentrator-5 kit (Zymo Research, USA). These amplified copies of the original DNA isolations contain no DNA modifications, and hence were used as a negative control.

Barcoded ONT libraries were built from each raw phage DNA isolation and WGA copy using the Rapid Barcoding Kit 96 RBK 110-96 (ONT, UK). The libraries were then sequenced together on the MinION platform using an R9 flowcell and MinKNOW.

### 3.4 Genomic characterization of nicked phages

To assemble the isolated phage genomes, Illumina reads were trimmed and quality-controlled using TrimGalore (v0.6.6)^23^ and assembled with SPAdes (3.13.1)^24^. As in previous work^18^, assemblies were manually quality-controlled and reoriented with respect to their correct start sites in CLC Genomics Workbench (v22). Phage genomes were annotated using Pharokka (v1.4.1)^25^. Similar phages were found using a combination of whole-genome BLASTN and tBLASTx searches (2.14.1+) against the INPHARED database (August 2023)^26^. In the case of Pseudomonas phage Rembedalsseter, a BLASTN search (webserver, October 2023) was also conducted to identify similar sequences in bacterial assemblies. Genomad (v1.7.0)^27^ was used to predict prophages in the bacterial assemblies with the best BLASTN hits, and the relevant prophage regions extracted.

For each phage isolate, selections of the top BLAST hits were made for further analysis. In each case, the top five BLASTN hits (ranked by score) were included in addition to a manual selection of other related phages. To quantify the intergenomic similarity between related phages, pairwise whole-genome nucleotide identities were calculated and visualized using VIRIDIC (webserver)^28^, and the recommended 70% genome identity cutoff from VIRIDIC was used to classify phages at the genus level^28,29^. Selected related phages were also visualized using clinker (v0.0.28)^30^ to show amino acid alignments between phage coding sequences. BACPHLIP (v0.9.6)^31^ was also used to predict the lifestyle (virulent/temperate) of each phage isolate. Phages with a virulent score > 0.5 were classified as virulent.

Using the Pharokka annotations, large terminase subunit (TerL) proteins were also identified, and a BLASTP (2.14.1+) search was conducted to identify the closest hits. Muscle (v3.8.1551)^32^ was used to align the TerL amino acid sequences, and IQTREE (v2.2.5)^33^ was used to generate nonparametrically bootstrapped (n=1000) phylogenetic maximum likelihood trees of the closest five BLASTP hits (ranked by score) along with a manual selection of other related phages. The trees were then visualized with ITOL v6^34^ and rooted at midpoint.

### 3.5 Environmental distribution of similar phages

To summarize the environments in which similar phages are found, we combined metadata from three sources; genus-level (70% nucleotide identity) phage isolates, genus-level metagenomically assembled contigs from the IMG/VR v4 database, and Sequence Read Archives (SRA) from metagenomic samples. Using Sourmash Branchwater^35^ to search for similar phages in >1,000,000 metagenomic SRAs, we reported SRAs which contain > 70% of the sampled phage 21-mers. We then combined the environmental metadata from these three sources to form a profile for each phage. We also downloaded reads from a single SRA (SRR14611561) with 100% containment to phage Kaldavass and mapped them to the Kaldavass genome using bwa-mem^36^ to verify read coverage.

### 3.6 Analysis of anti-defense systems and DNA modifications

The dbAPIS database consists of experimentally verified anti-defense proteins together with bioinformatically-derived protein families^37^. Using the derived protein families, we conducted an HMM search for similar proteins in the phage genomes, recording hits with an e-value < 0.05. For each hit, we subsequently conducted a BLASTP against the experimentally verified protein in each family.

To detect phage genome nucleotide modifications in the Nanopore sequencing data, we used the TOMBO_HP workflow^38^. In brief, raw sequencing signals were basecalled using Guppy 6.3.9 with the high-accuracy model ’dna_r9.4.1_450bps_sup.cfn’. The basecalled reads were aligned to the phage genomes using minimap2 (v2.24)^39^. Then, the raw signal from the mapped reads was anchored to the reference genome (resquiggled) using Tombo (1.5.1). This software was used to detect modifications, by comparing the signal from the native DNA to that of the control sample that underwent Whole Genome Amplification (WGA). The modified fraction (modfrac) statistics from Tombo were used to generate plots and tables that were manually inspected to detect the DNA modification motifs. Known motifs associated with Dam and Dcm methylation were further examined with Tombo using motif-specific models available for *Escherichia coli* Dam and Dcm methylation to confirm the presence of these specific methylation events.

For the motif analysis of Pseudomonas phage Rembedalsseter, the 250 closest phage genomes were selected by tBLASTx search against the INPHARED database (August 2023), ranked by aggregated score per genome. Distances between all genome pairs were calculated using the weighted gene repertoire relatedeness (wGRR)^40^. Occurrences of the motif [TA]ACT[GA]TGAC were located in each genome using seqkit v2.5.1^41^. Pharokka was used to gene-call the 60 closest genomes and reorientate them relative to the conserved *terL* gene. Motif positions were then calculated relative to genome length and coding sequences.

## 4. Results and Discussion

Eight phages were isolated from organic waste samples using seven bacterial strains previously isolated from the wheat phyllosphere^15^ as replication hosts: four *E. aphidicola* strains, two *P. poae* strains, and one *P. trivialis* strain. Host genome assemblies available under NCBI BioProject PRJNA951732.

### Erwinia phages Vettismorki and Gravdalen

The two closely related Erwinia phages Vettismorki and Gravdalen were isolated against the *E. aphidicola* strains B01_5 and N2_6, respectively.

TEM imaging showed Erwinia phages Vettismorki and Gravdalen to be tailed, contractile phages. Vettismorki had a tail length of 119±4 nm, head length of 121±4 nm, and head width of 90±2 nm, while Gravdalen had a tail length of 117±2 nm, head length of 119±4 nm, and head width of 85±3 nm. Both phages also have long tail fibers (shown for Gravdalen in Fig. 1B). Vettismorki and Gravdalen are very similar genetically (93.5% nucleotide similarity), with genome sizes of 171,806 bp and 171,558 bp and GC contents of 38.62% and 38.66%, respectively.

**Figure 1.**
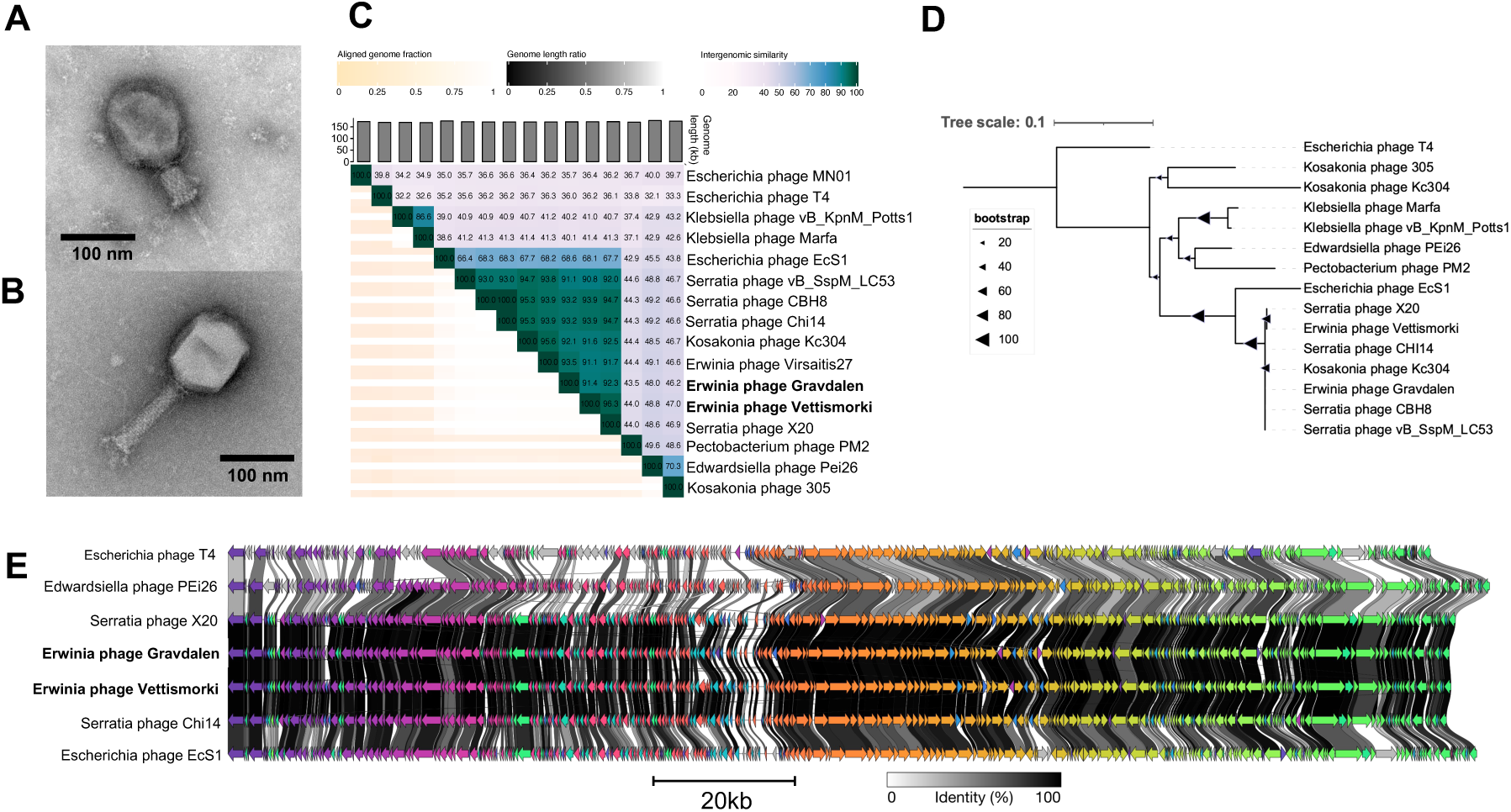
Analysis of Erwinia phages Vettismorki and Gravdalen. **A)** TEM imaging of Erwinia phage Vettismorki, tail contracted. **B)** TEM imaging of Erwinia phage Gravdalen, tail non-contracted and tail fiber visible. **C)** Whole-genome nucleotide similarity of Vettismorki and Gravdalen in comparison with select related phage genomes (VIRIDIC). **D)** Bootstrapped maximum likelihood tree of large terminase subunit amino acid alignments with select related phages. **E)** clinker figure showing amino-acid similarities between protein-coding sequences of select related phages.

Phylogenetically, Erwinia phages Vettismorki and Gravdalen are closest to Serratia phages X20 and Chi14^42^ and Erwinia phage Virsaitis27, both with respect to whole genome nucleotide identity and the TerL maximum likelihood tree (Figs. 1C-D). All three of these phages are classified within the genus *Winklervirus*, therefore also placing Vettismorki and Gravdalen within *Winklervirus*. Vettismorki is so closely related to X20 (96.3% nucleotide similarity) that it should be placed in the same species; Winklervirus xtwenty. The genus *Winklervirus* lies within the subfamily *Tevenvirinae*, named for the representative species Escherichia phage T4, and Vettismorki and Gravdalen are shown to be related to T4 (Fig. 1C-E).

### Erwinia phages Hallingskeid and Kaldavass

Erwinia phages Hallingskeid and Kaldavass were isolated against *E. aphidicola* strains B01_10 and W09_2, respectively.

TEM imaging showed both phages are large phages with contractile tails. Hallingskeid had a tail length of 131±4 nm, head length of 136±3 nm, and head width of 111±8 nm, while Kalvdavass had a tail length of 128±5 nm, head length of 131±3 nm, and head width of 112±6 nm. Both phages also have short tail fibers terminating in rounded appendages almost resembling small feet. Related jumbo phages (Pectobacterium phage CBB, Klebsiella phage Rak2) are known to encode complex tail fiber assemblies^43,44^ and these feet-like appendages may be analogous complexes. Interestingly, both Hallingskeid and Kaldavass have multiple proteins annotated as tail fibers (two and six proteins, respectively), further supporting this interpretation.

Sequencing showed that the large capsids of the viral particles were reflected in genome size; Hallingskeid and Kaldavass were 366,158 bp and 366,556 bp respectively, far above the 200 kbp cutoff for jumbo phages. Although the two phages are of similar size and morphology, they share little nucleotide identity (26.3% whole-genome similarity). They are also related to other Enterobacterial jumbo phages of similar sizes (Fig. 2C). In particular, Kaldavass is similar to Xanthomonas phage XbC2 (84.6% similarity) with related TerL proteins(Fig. 2D). Meanwhile, Hallingskeid is somewhat similar to Escherichia phage EJP2^45^ (39.5%) and has a more diverged TerL. Hence, Kaldavass should be placed in the genus *Mimasvirus* with XbC2, while Hallingskeid should form a novel genus. More broadly, all the aforementioned phages reside within the so-called group 2.2 jumbo phages which are distant descendants of the smaller *Tevenvirinae* phages^46^.

**Figure 2.**
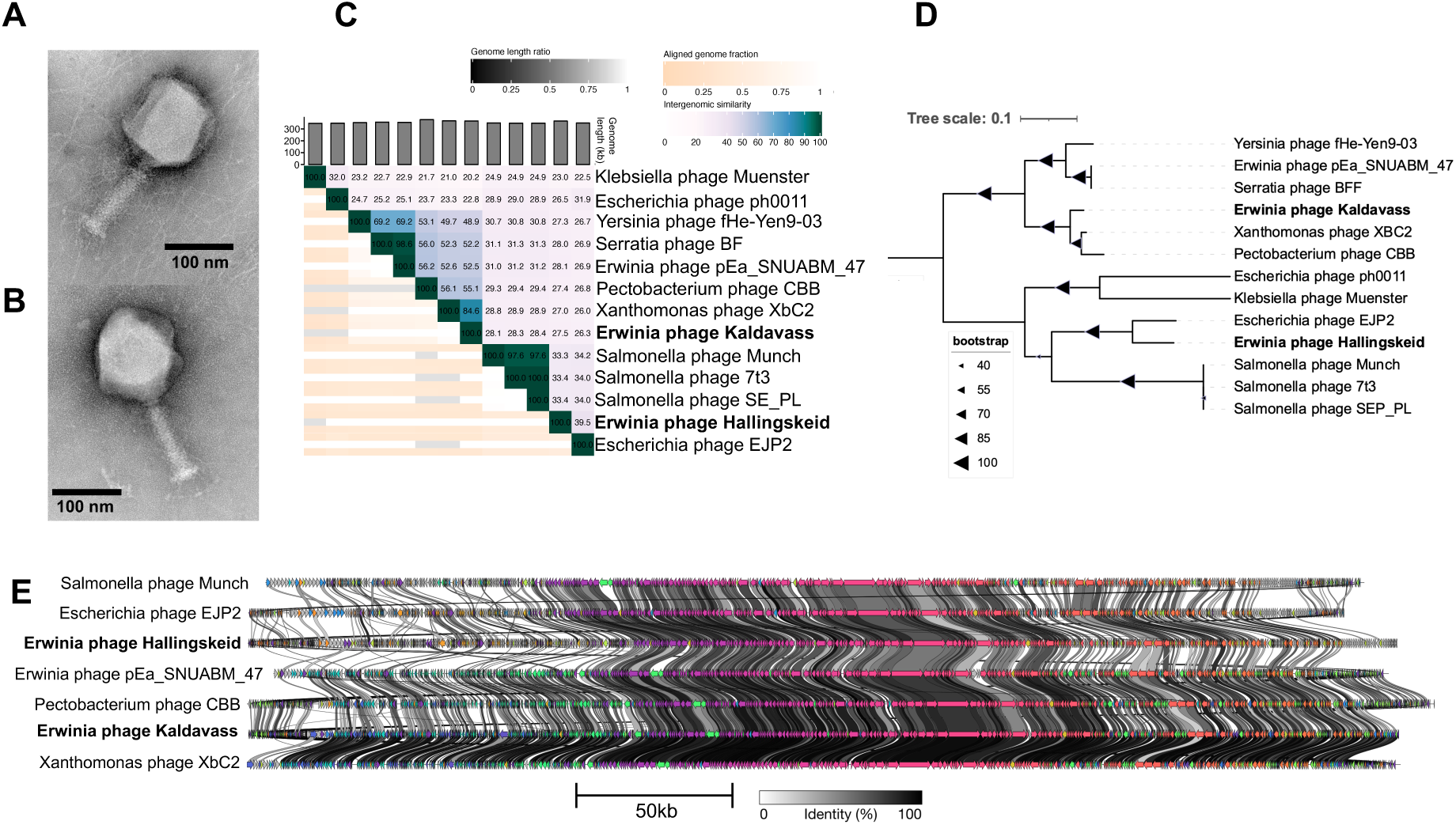
Analysis of Erwinia phages Hallingskeid and Kaldavass. **A)** TEM imaging of Erwinia phage Hallingskeid, with tail non-contracted and tail fibers visible. **B)** TEM imaging of Erwinia phage Kaldavass, with tail non-contracted and tail fibers visible. **C)** Whole-genome nucleotide similarity of Hallingskeid and Kaldavass in comparison with select related phage genomes (VIRIDIC). **D)** Bootstrapped maximum likelihood tree of large terminase subunit amino acid alignments with select related phages. **E)** clinker figure showing amino-acid similarities between protein-coding sequences.

Although this group of phages diverge at the nucleotide level, aligning their proteins reveal a common architecture (Fig. 2E). Roughly, the genomes 150 kbp conserved region consisting primarily of well- annotated DNA/RNA metabolism and structural genes, flanked by variable 100 kbp regions on either side with many smaller genes of unknown functions.

### Pseudomonas phage Rembedalsseter

Pseudomonas phage Rembedalsseter was isolated on the *P. trivialis* strain B08_4.

TEM imaging shows this phage to have a symmetric icosahedral head (length and width 62±4 nm and 60±3 nm) and a tail length of 40±6 nm (Fig. 3A). Rembedalsseter’s genome reflects this modest size (45,228 bp) and has a GC content of 52.5% (Table 1). The closest whole-genome hits to Rembedalsseter are Pseudomonas phages UFV-P2^47^ and NV1^48^ (76.7% and 69.5% whole-genome similarity respectively, Fig. 3B). TerL alignments also show these two phages to be Rembedalsseter’s closest neighbors (Fig. 3C). As both these phages reside within the genus *Vicosavirus,* Rembedalsseter should accordingly be placed within this genus. *Vicosavirus* is closely related to the genus *Bruynoghevirus*, which includes the well-studied Pseudomonas phage LUZ24^49^. Members of both genera share considerable synteny (Fig. 3D), with bidirectionally transcribed DNA coding for early and middle genes on the left, and late genes on the right.

**Figure 3.**
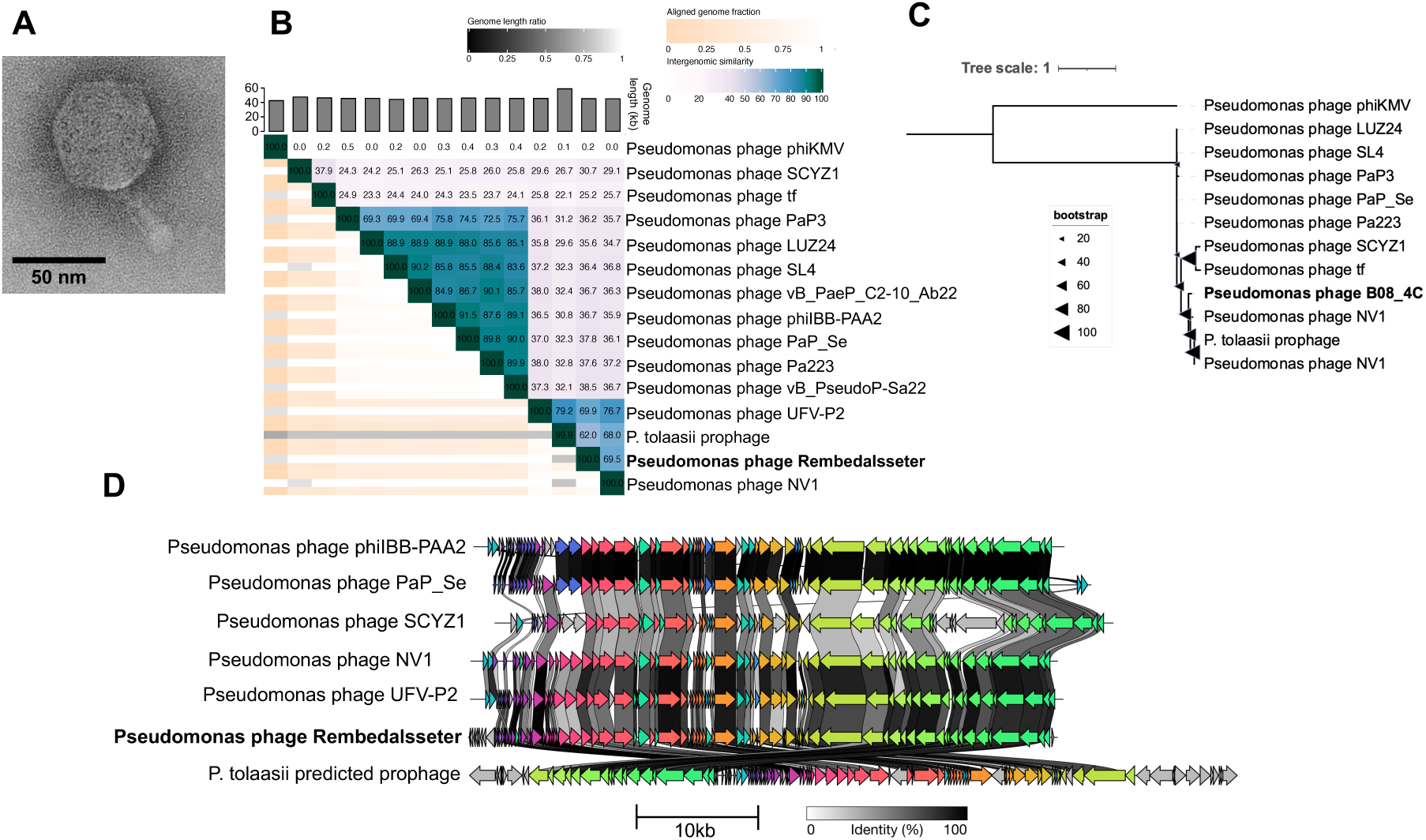
Analysis of Pseudomonas phage Rembedalsseter. **A)** TEM imaging of Pseudomonas phage Rembedalsseter. **B)** Whole-genome nucleotide similarity of Rembedalsseter in comparison with select related phage genomes (VIRIDIC). **C)** Bootstrapped maximum likelihood tree of large terminase subunit amino acid alignments with select related phages. **D)** Clinker figure showing amino-acid similarities between protein-coding sequences. The *P. tolaasii* prophage prediction is in the same orientation as it appears in the bacterial genome.

**Table 1.** Overview of the 8 isolated phages, showing isolation host, genome length, GC content, and proposed genera. All phages are isolated from household organic waste from Glostrup, Denmark. * Pseudomonas phage Milchi should be placed into the same genus as Pseudomonas phage Astolliot, but this phage is currently misclassified within an incorrect genus (*Otagovirus*).

| Name | Host | Length (bp) | GC (%) | Proposed genus |
| --- | --- | --- | --- | --- |
| Erwinia phage Vettismorki | *E. aphidicola* B01_5 | 171,806 | 38.6 | *Winklervirus* |
| Erwinia phage Gravdalen | *E. aphidicola* N2_6 | 171,558 | 38.7 | *Winklervirus* |
| Erwinia phage Hallingskeid | *E. aphidicola* B01_10 | 366,158 | 39.5 | *Mimasvirus* |
| Erwinia phage Kaldavass | *E. aphidicola* W09_2 | 366,556 | 38.4 | Novel genus |
| *Pseudomonas* phage Rembedalsseter | *P. trivialis* B08_4 | 45,228 | 52.5 | *Vicosavirus* |
| *Pseudomonas* phage Boevelstad | *P. poae* B04_10 | 92,884 | 44.9 | Novel genus |
| *Pseudomonas* phage Torfinnsbu | *P. poae* Z9_5 | 53,022 | 56.9 | Novel genus |
| *Pseudomonas* phage Milchi | *P. poae* Z9_5 | 174,859 | 45.5 | Novel genus* |

**Table 2.** HMM search for known anti-defense systems in the eight phage genomes using the dbAPIS database. Hits to APIS protein families were followed up with a BLASTP against the experimentally verified seed protein from each protein family. The bacterial defense system targeted by each verified anti-defense system is also shown.

| Phage | APIS family | HMM e-value | Experimentally verified seed | BLASTP ID to seed | BLASTP query cover to seed | Targeted bacterial defense system |
| --- | --- | --- | --- | --- | --- | --- |
| Rembedalsseter | AcrIF7 | 1.6E-03 | AcrIF7 | 0.36 | 0.08 | CRISPR |
| Milchi | APIS009 | 2.7E-04 | Ral | 0.19 | 0.41 | RM |
| Milchi | APIS068 | 3.2E-03 | Gad1 | 0.5 | 0.11 | Gabija |
| Milchi | APIS055 | 1.1E-03 | KlcA | 0 | 0 | RM |
| Milchi | APIS016 | 2.3E-28 | Acb1 | 0.44 | 0.89 | CBASS |
| Milchi | APIS023 | 5.3E-05 | gp5.9 | 0.3 | 0.3 | RecBCD |
| Milchi | APIS060 | 7.5E-07 | Apyc1 | 0 | 0 | Pycsar |
| Milchi | APIS063 | 3.4E-06 | ArdA | 0.25 | 0.26 | RM |

Interestingly, these *Vicosavirus* phages share considerable (>60%) whole-genome similarity) with a predicted prophage from bacterial strain *P. tolaasii* FP2293. In fact, UFV-P2’s TerL is 100% identical to that of the *P. tolaasii* prophage, indicating a very close phylogenetic relationship. Similarly, the more-distantly related *Bruynoghevirus* PaP3 was shown to integrate within its bacterial host’s chromosome despite lacking any detectable integrase or transposase^50^. However, phage lifestyle predictor BACPHLIP predicts all *Vicosavirus* and *Bruynoghevirus* members to be virulent, and no integrases or transposases were found in any of the phages (including the *P. tolaasii* prophage) using Pharokka. At this point, it is unclear how PaP3 or the *P. tolaasii* prophage establish lysogeny, or whether the other *Vicosavirus* and *Bruynoghevirus* phages may have a temperate lifestyle^51^.

### Pseudomonas phage Boevelstad

Pseudomonas phage Boevelstad was isolated from plaques on host strain *P. poae* B04_10.

Pseudomonas phage Boevelstad has a symmetrical icosahedral head (length and width 74±3 nm and 72±2 nm respectively), 132±6 nm contractile tail, and visible tail fibers (Fig. 4A). Although Boevelstad’s tail is roughly the same size as those of jumbo phages Hallingskeid and Kaldavass, the head is much smaller, perhaps reflecting Boevelstad’s smaller genome of 92,884 bp. Genetically, Boevelstad is quite distant from all NCBI phage genomes besides Pseudomonas phage PseuGes_254, at 65.6% genome similarity (Fig. 4B). The TerL maximum likelihood tree also shows close relation to PseuGes_254, which has not yet been placed in a named genus (Fig. 4C). However, Boevelstad is distinct enough to be placed in a new genus separate from PseuGes_254.

**Figure 4.**
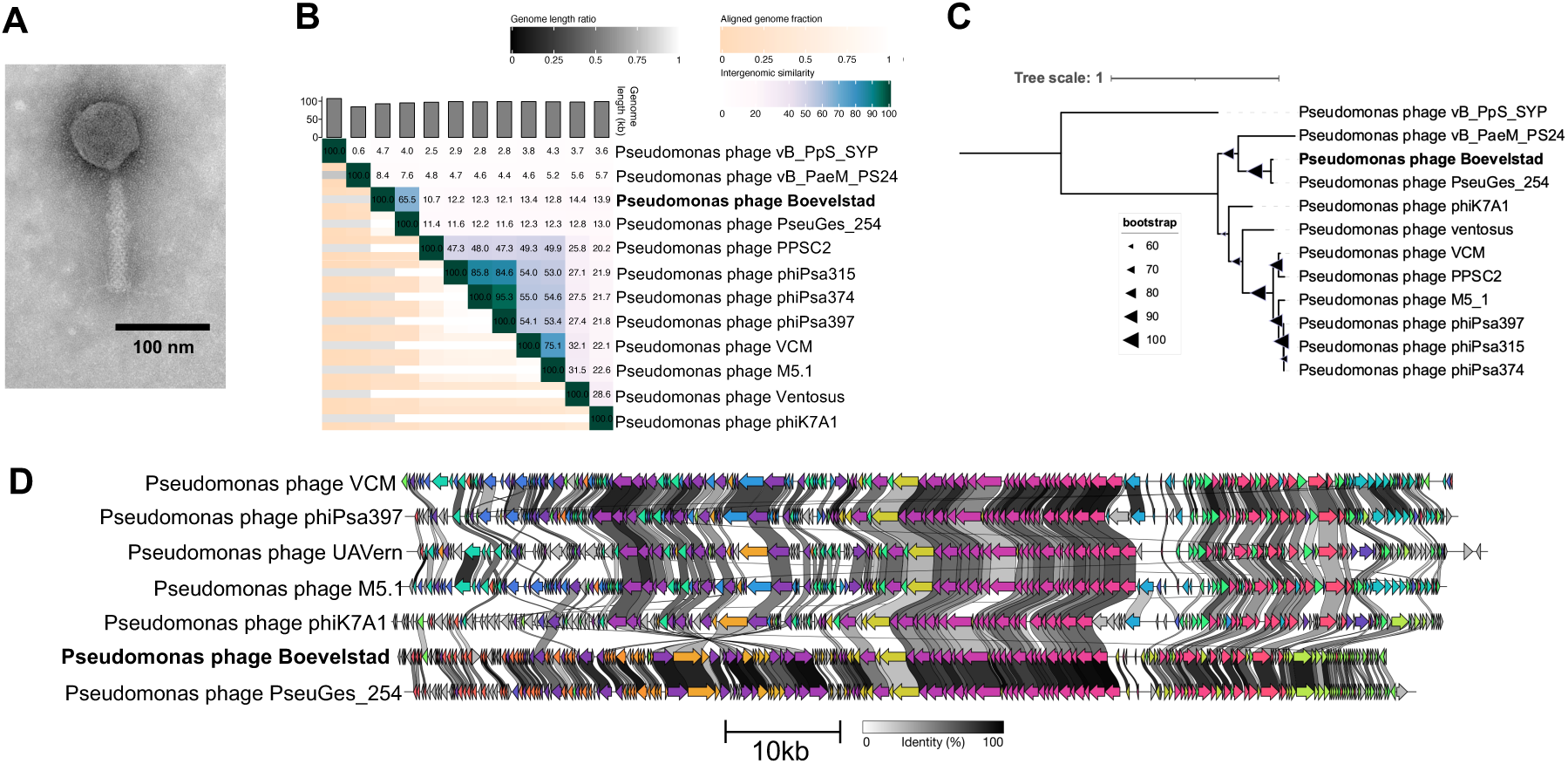
Analysis of Pseudomonas phage Boevelstad. **A)** TEM imaging of Pseudomonas phage Boevelstad. **B)**Whole-genome nucleotide similarity of Boevelstad in comparison with select related phage genomes (VIRIDIC). **C)** Bootstrapped maximum likelihood tree of large terminase subunit amino acid alignments with select related phages. **D)** Clinker figure showing amino-acid similarities between protein-coding sequences.

Genome alignment also shows close relation between Boevelstad and PseuGes_254, although there is also substantial alignment with other Pseudomonas phages such as phiK7A1 and M5.1 (Fig. 4D). Relative to the other phages shown in Fig. 4D, Boevelstad and PseuGes_254 have a large inversion (bases 24,023 – 40,934 in Boevelstad). This genomic region codes for many proteins related to nucleotide metabolism, such as DNA primase/helicase, DNA polymerase I, and ribonucleotide reductase. Also of note is a predominantly non-protein-coding region (approx. 3.5 kbp, bases 66,796 - 70,162 in Boevelstad) present in all phages shown in Fig. 4D. This region is host to many tRNAs; in Boevelstad, 12 tRNAs are located here.

### Pseudomonas phage Torfinnsbu

Pseudomonas phage Torfinnsbu was isolated on the bacterial strain *P. poae* Z9_5.

Morphologically, Torfinnsbu has a symmetric icosahedral head (length and width 69±2 nm and 67±2 nm respectively) and a long (159±8 nm) flexible, striated, tail (Fig. 5A). Torfinnsbu has a 53,022 bp genome with a somewhat high GC content (56%). However, this is slightly below the GC content of the host strain *P. poae* Z9_5 (60%) as is normal for phages^52^.

**Figure 5.**
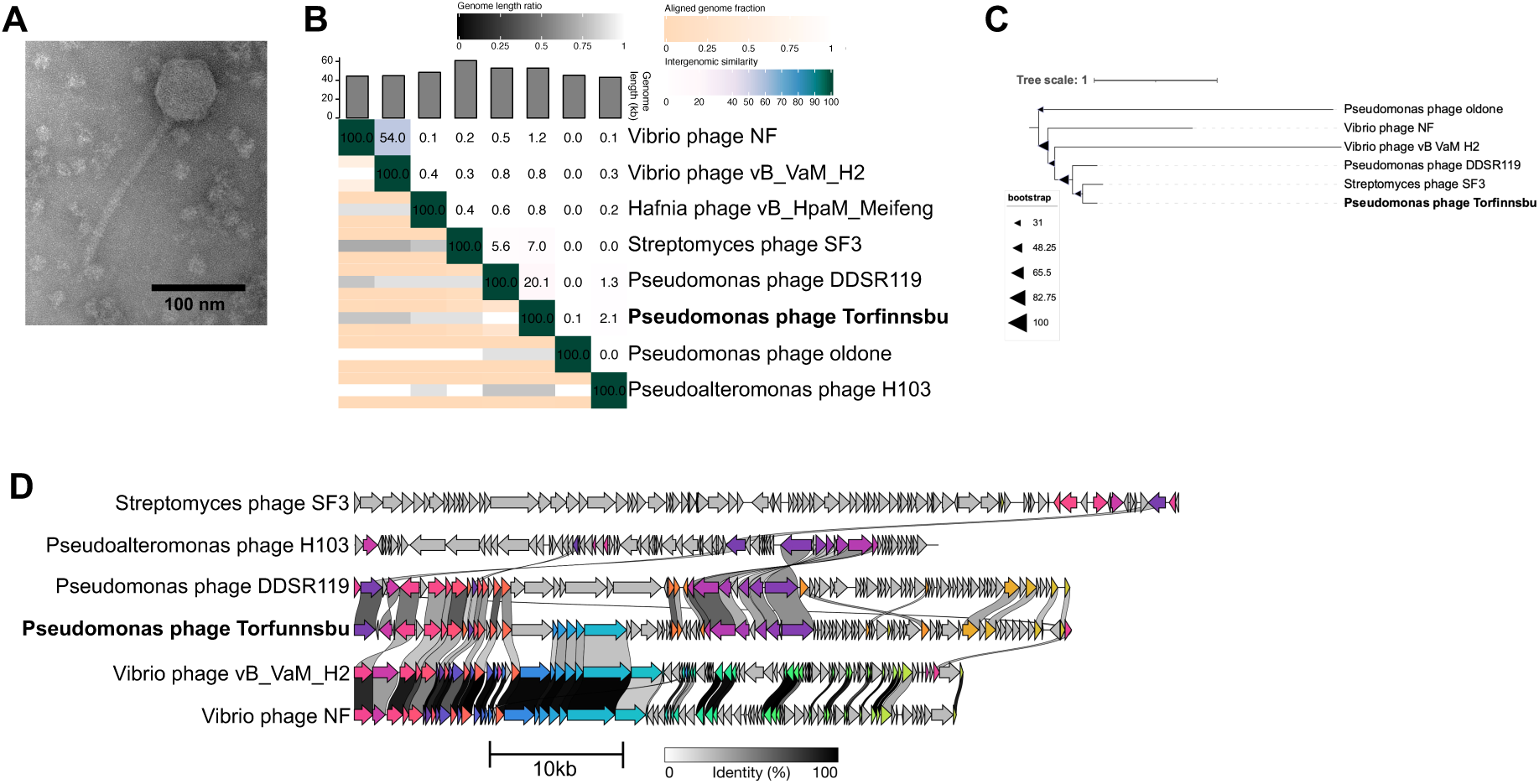
Analysis of Pseudomonas phage Torfinnsbu. **A)** TEM imaging of Pseudomonas phage Torfinnsbu. **B)**Whole-genome nucleotide similarity of Torfinnsbu in comparison with select related phage genomes (VIRIDIC). **C)** Bootstrapped maximum likelihood tree of large terminase subunit amino acid alignments with select related phages. **D)** Clinker figure showing amino-acid similarities between protein-coding sequences.

Phylogenetically, however, Torfinnsbu does not have any close relatives among known phage isolates. Pseudomonas phage DDSR119 has 20% whole-genome nucleotide similarity to Torfinnsbu, but no other phage isolates have any substantial nucleotide similarity (Fig. 5B). Although the maximum likelihood tree (Fig. 5C) shows Streptomyces phage SF’s TerL as the closest hit phylogenetically, the two phages otherwise share very few genes (Fig. 5D) and are therefore only very distantly related.

Whole-genome protein alignments (Fig. 5D) show Torfinnsbu shares homology with Vibrio phages vB_VaM_H2 and Vibrio phage NF over a leftmost region of ∼22 kbp coding for structural genes. Pseudomonas phage DDSR119 also shares amino acid similarity with some of Torfinnsbu’s structural genes, in addition to a downstream homologous region coding for DNA metabolism-associated proteins. Interestingly, one of Torfinnsbu’s proteins is annotated as a ParB-like partition protein, which might indicate some type of plasmid-phage lifecycle. Although no other annotations were found that relate to plasmid functions, this protein is also conserved in DDSR119.

### Pseudomonas phage Milchi

Pseudomonas phage Milchi was also isolated on the *P. poae* strain Z9_5.

Pseudomonas phage Milchi has a nearly symmetrical icosahedral head (length and width 133±3 nm and 119±7 nm) a 129±5 nm contractile, striated tail terminating in straight tail fibers (Fig. 6A). Milchi is a large phage with a genome length of 174,859 bp and encodes 29 predicted tRNAs.

**Figure 6.**
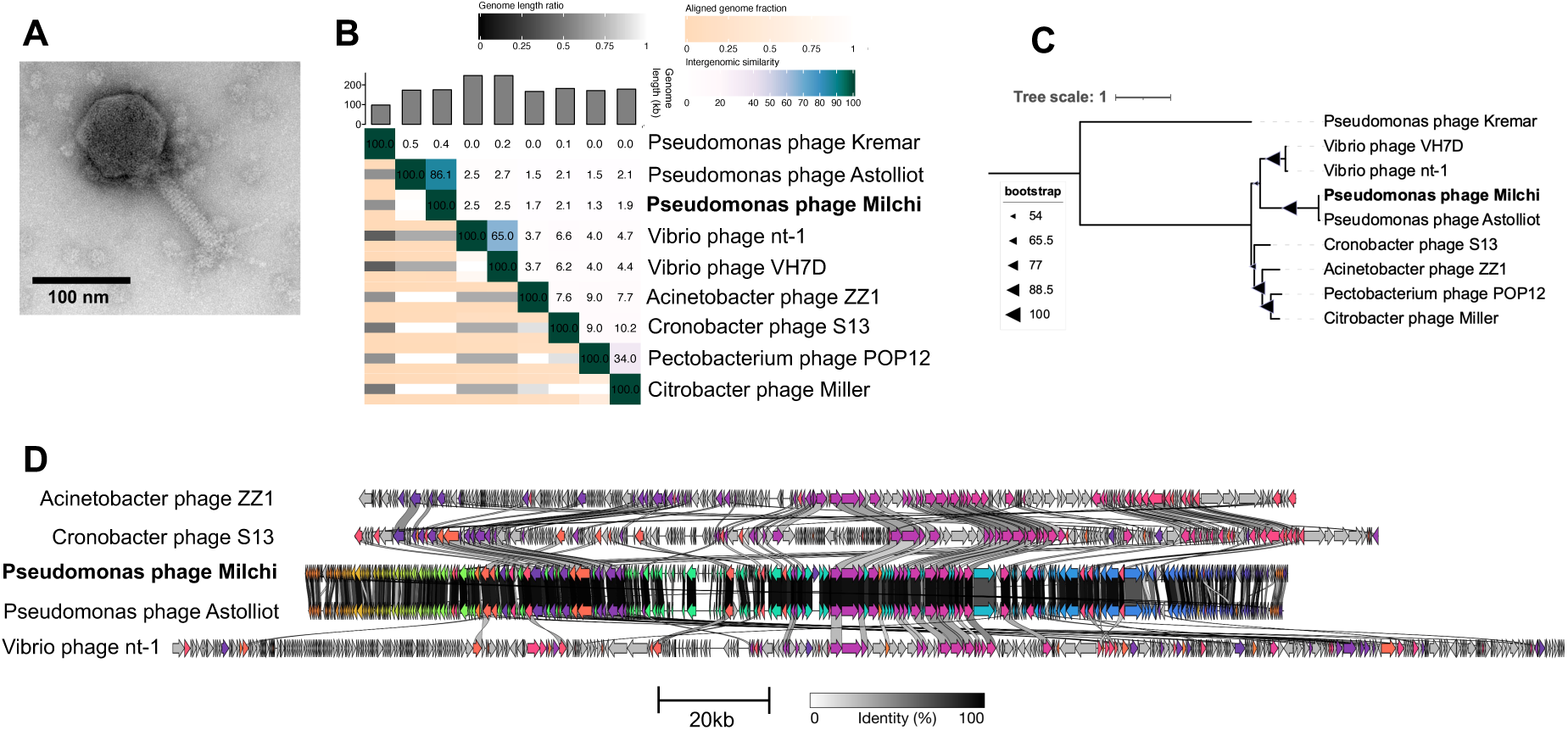
Analysis of Pseudomonas phage Milchi. **A)** TEM imaging of Pseudomonas phage Milchi. **B)** Whole-genome nucleotide similarity of Milchi in comparison with select related phage genomes (VIRIDIC). **C)** Bootstrapped maximum likelihood tree of large terminase subunit amino acid alignments with select related phages. **D)** Clinker figure showing amino-acid similarities between protein-coding sequences

Phylogenetically, Milchi is closely related to Pseudomonas phage Astolliot, with a whole genome nucleotide similarity of 86.1% (Fig. 6B), closely related TerL proteins (Fig. 6C) and highly conserved synteny and coding sequences (Fig 6D). Based on this, Milchi should be placed in the same genus as Astolliot. On NCBI, Pseudomonas phage Astolliot has been placed into the genus *Otagovirus*, which has two ICTV-recognized species: Pseudomonas phage VCM and Pseudomonas phage phiPsa374^53^. However, a VIRIDIC nucleotide comparison found that neither Milchi nor Astolliot has any significant nucleotide identity (<0.2%) with members from either species. Using the commonly accepted 70% nucleotide identity threshold for phage genera, Milchi and Astolliot should therefore be reclassified and placed in a novel, separate genus as two new species.

### Environmental distribution of similar phages

As all eight of these phage isolates were obtained from a different environment (organic waste) than their bacterial hosts (wheat flag leaf), we were interested in which other environmental niches these phages inhabit. To investigate this, we combined environmental metadata from similar (>0.7 nucleotide similarity) phage isolates, metagenomically assembled phage contigs, and metagenomic SRAs. To search SRAs directly for similar phages, we used Branchwater^35^ which samples a set of 21-mers from the query genome and compares them to sampled 21-mers from > 1,000,000 publicly available SRAs. We included hits with >70% kmer containment as an approximation of genus-level hits. Combining hits from isolates, metagenomic assemblies, and SRA, we analyzed the sample isolation environments (Supplementary table 1).

No phages were found with genus-level similarity to Pseudomonas phages Boevelstad or Torfinnsbu in publically available isolate databases or metagenomes. This underlines both the need for phage isolations, the novelty of these two phages, and the sheer diversity of phage genomes worldwide; in > 1,000,000 metagenomic SRAs, no genus-level hits were found. In contrast, genus-level hits were found for Pseudomonas phages Milchi and Rembedalsseter, and Erwinia phages Vettismorki, Gravdalen, Hallingskeid, and Kaldavass (Fig. 7A). Interestingly, only the jumbo phages Hallingskeid and Kaldavass had metagenomic hits (IMGVR or Branchwater); all seven Hallingskeid hits were metagenomic, while 44/45 Kaldavass hits were metagenomic. This could be due to biases in phage isolation; for example, the crAssphage family is the most abundant in human gut microbiomes, yet the first crAss-like phage isolate was reported in 2018^54^.

**Figure 7.**
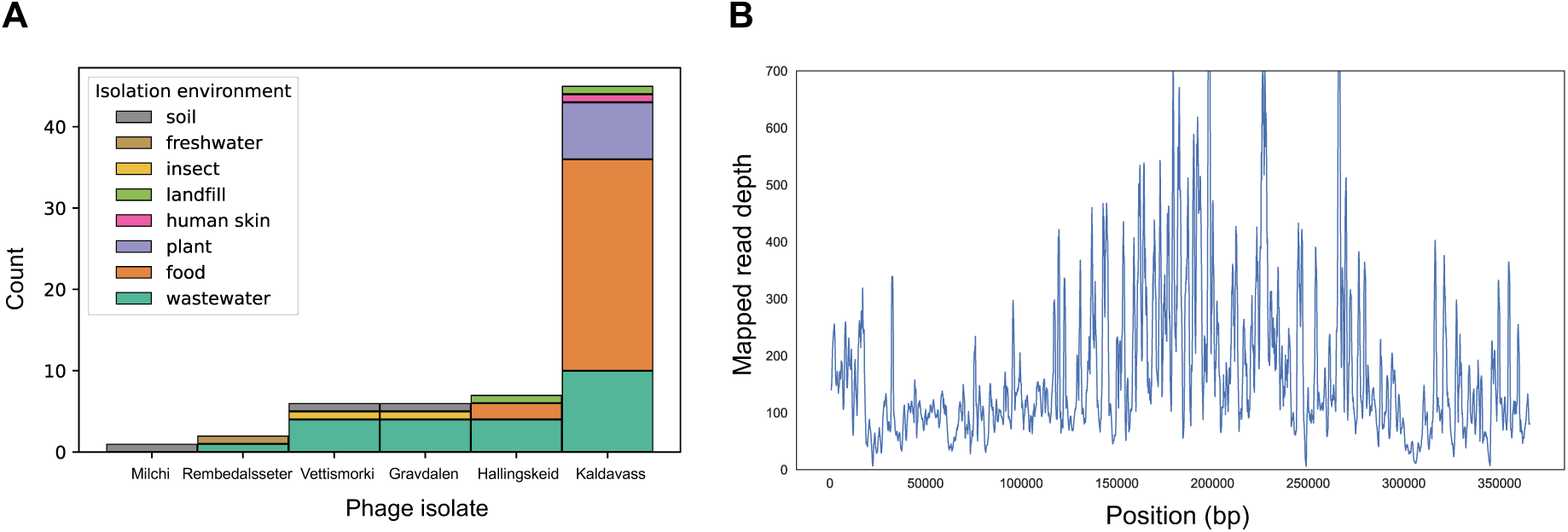
**A)** Breakdown of environments where similar phages were found. Environmental metadata was combined from similar (>70% nucleotide similarity) phage isolates and metagenomically assembled contigs, and SRAs with > 70% 21-mer containment, determined using Branchwater. **B)** Read mapping from a Spanish spinach virome (SRR14611561) to the genome of phage Kaldavass, demonstrating near-complete genome coverage. Rolling average with window size = 1000 bp.

Although all phages were isolated from organic waste, genus-level phage hits were found in a greater variety of environments. Five of the phage isolates had hits from wastewater, illustrating why wastewater is so commonly used as a source for phage isolates. However, phage Kaldavass (45 hits) was dominated by hits to food and plant metagenomes. In each case where the food item was specified, it was a plant-based food, underlining how this phage appears to be conserved in a variety of plant microbiomes. Astoundingly, one Spanish spinach virome (SRR14611561^55^) had 100% kmer containment at 100% identity to Kaldavass, indicating the presence of a (near) identical phage. To confirm this, we mapped the reads from this sample to Kaldavass and found only 227/366,556 bases in the entire genome (0.06%) had zero coverage (Fig. 7B). More broadly, the fact that a jumbo phage from organic waste in Denmark is near-identical to a phage from spinach in Spain and closely related to phages in American wastewater, speaks to how interconnected microbiomes are.

### Phage-encoded anti-defense systems and DNA modifications

Next, we searched for anti-defense systems and DNA modifications in the eight phage genomes. Using the dbAPIS^37^ database of protein families derived from experimentally verified anti-defense systems, we conducted an HMM search for similar proteins in the phage genomes, followed by a BLASTP against the experimentally verified protein seed in each family.

Only two of the phages had HMM hits to the dbAPIS database; Pseudomonas phages Rembedalsseter and Milchi. Interestingly, Milchi had 7/8 total dbAPIS hits from the nine phages, suggesting this phage may carry an especially high number of antidefense systems. However, these are very distant hits and should be treated with caution. Except for one good (89% coverage, 44% identity) Milchi hit to the anti-CBASS protein Acb1, BLASTP query coverage and identity were very low, and these proteins may not play any role in anti-defense. Conversely, we should not conclude that the other seven phages do not encode anti-defense systems. With dbAPIS only containing 41 experimentally verified seed proteins, we have only begun to scratch the surface of the phage repertoire of anti-defense systems.

In addition to explicit anti-defense systems, phages commonly use DNA modifications to avoid recognition by bacterial anti-phage defense systems such as RM and CRISPR-Cas9^22,56,57^. We therefore searched for DNA modifications using Nanopore sequencing. By comparing the electrical signals generated from native phage DNA with that of WGA genome copies, we detected motifs associated with shifts in raw current signal levels, indicative of DNA modifications^13^. These motifs, along with their predicted modification type, are shown in Table 3.

**Table 3.** Summary of proposed DNA modifications, phage-encoded DNA methyltransferases, and modification motifs for each of the eight phages. Bold font indicates the modified base in the motif, while brackets indicate multiple variable bases. Known methylation motifs are indicated with *. Ara-hmc is arabinose-hydroxymethylcytosine, a modification hypothesized based on protein homology with Escherichia phage RB69.

| Phage | # DNA methyltransferases | Proposed modification type | Motifs |
| --- | --- | --- | --- |
| Vettismorki | 0 | ara-hmc | unknown |
| Gravdalen | 0 | ara-hmc | unknown |
| Hallingskeid | 2 | methylation | GATC*, CC[AT]GG* |
| Kaldavass | 3 | methylation + unknown | GATC*, CC[AT]GG*, TGCAT |
| Rembedalsseter | 0 | nicked DNA | [AT]ACT[AG]TGAC |
| Boevelstad | 1 | None |  |
| Torfinnsbu | 0 | None |  |
| Milchi | 0 | None |  |

In contrast to the six other phages, native DNA from Erwinia phages Vettismorki and Gravdalen could not be sequenced with Nanopore technology. Although the WGA genomes could be sequenced and mapped, the native DNA reads were of so low quantity and quality that they could not be mapped back to their reference genomes. This is a known phenomenon when sequencing T-even phages with Nanopore technology, many of whom are known to be hypermodified with sugar molecules throughout the entire genome that greatly reduce read quality (shown for glucosyl-hydroxymethylcytosine (glc-hmC) in T4, and arabinose-hydroxymethylcytosine (ara-hmC) in RB69)^13^. In this case, Vettismorki and Gravdalen have proteins with (distant) homology to both T4-encoded dCMP hydroxymethyltransferase and dNMP kinase which produce hmC in T4^58^. However, the Erwinia phages have no homologs to the alpha and beta-glucosyltransferases required to produce glc-hmC in T4, or the proposed arabinosyltransferase found in RB69^59^. They do however have close homologs to the RB69-encoded ORF53_52C and ORD55C which are proposed to be involved in the synthesis of UDP-arabinose, a precursor to the ara-hmc^59^ modification. However, additional analysis is required to confirm the identity of the proposed hypermodification.

Of the eight phages investigated, two (Erwinia phages Hallingskeid and Kaldavass) displayed probable DNA modifications at two well-known methylation motifs; GATC and CC[AT]GG. These motifs are methylated in many bacteria and phages (modified by Dam and Dcm, respectively, in *E. coli*)^60^. We further analysed the modification data using motif-specific Tombo models for *E. coli* Dam and Dcm, confirming the presence of 6-methyladenine at GATC and 5-methylcytosine at CC[AT]GG. Erwinia phage Kaldavass was also modified at an additional motif (TGCAT). However, this motif is not present in the REBASE databases of restriction recognition systems and PacBio-identified methylation motifs^61^ and hence cannot be immediately identified. Each of the three methylated phages also encoded its own DNA methyltransferases, although there was not a one-to-one correlation between the number of encoded methyltransferases and the number of modification motifs; Pseudomonas phage Boevelstad also encodes a predicted DNA methyltransferase, but did not have any detectable modification motifs.

### Pseudomonas phage Rembedalsseter has 13 single-stranded DNA breaks associated with a motif conserved in a large family of Pseudomonas phages

No DNA modification motifs were found in Pseudomonas phage Rembedalsseter. However, when investigating the native single-stranded sequencing depths, we were surprised to see that the sequencing depth on the negative strand abruptly fell to zero (or near-zero) at 13 places throughout the genome (Fig. 8A). These coverage drops were not observed in the native positive strand, or in either strand of the WGA control. Analysing these 13 regions with MEME revealed a common motif present at all 13 sites: [AT]ACT[AG]TGAC. This motif occurred precisely 13 times, all on the same strand, and perfectly correlating with the coverage drops.

**Fig. 8.**
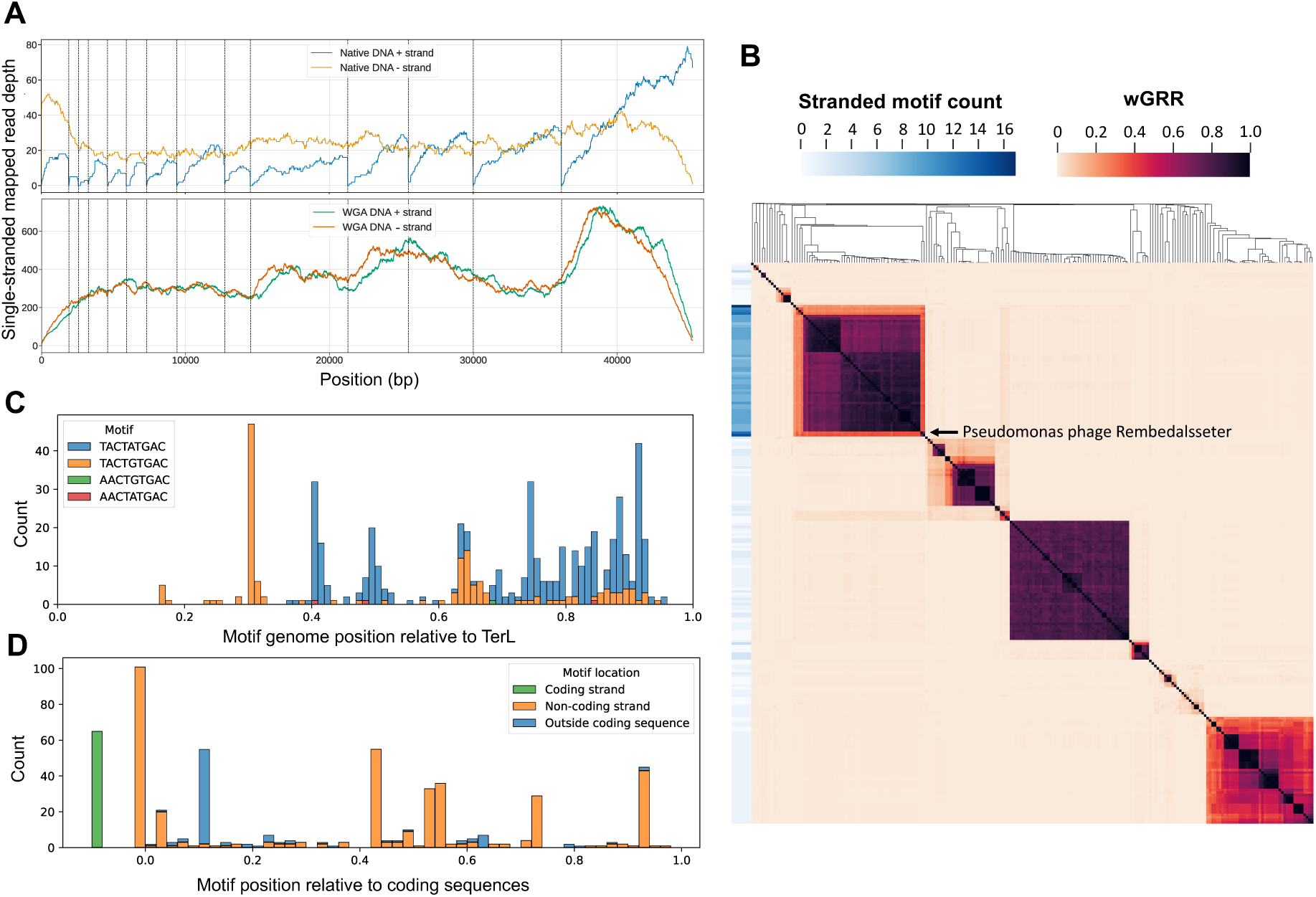
Investigation of motifs associated with the single-stranded DNA breaks in Pseudomonas phage Rembedalsseter and comparison to related phages. **A)** Single-stranded read-mapping depths of native and WGA DNA from Rembedalsseter. Dashed lines indicate [AT]ACT[GA]TGAC motifs. **B)** Heatmap of wGRR between Rembedalsseter and related phages with clustered rows and columns. Row labels indicate the difference in the number of [AT]ACT[GA]TGAC motif occurrences between the two strands of each phage genome. Pseudomonas phage Rembedalsseter is indicated. **C)** Position of motif locations relative to complete genome length in Rembedalsseter and the 60 related phages, with all genomes reoriented to start from *terL*. Bar colors indicate motif identity. **D)** Motif locations relative to their position to the coding sequences they occur within (0 is the start of a coding sequence, and 1 is the end). Bar colors indicate motif strand relative to the coding sequence, and whether or not it occurs outside a coding sequence.

A very similar motif (TACT[AG]TG[AC]C) was previously found to be associated with single-stranded DNA breaks (nicks) in the related Pseudomonas phage tf^62^ (relationship shown in Fig. 3B-C). Single-stranded DNA breaks explain the Nanopore sequencing depth pattern; on one strand, no reads span the motif, and hence the coverage drops to zero at these motifs. Single-stranded breaks have also been observed in several other phages, including the distantly related Pseudomonas phages phiKf77 and phiKMV^63^ (with a different motif), as well as the completely unrelated Escherichia phage T5^64^. In all cases, the function of these nicks is unclear. Nick-less T5 mutants have been isolated, indicating the nicks are not necessary for T5 growth under laboratory conditions. Several explanations have been proposed, including the facilitation of DNA packaging^62^ and gene expression^63^.

To investigate how widespread Rembedalsseter’s nick-associated motif ([AT]ACT[GA]TGAC) is, we searched the INPHARED database for phages with similarity to Rembedalsseter and gathered the 250 phages with the highest aggregate tBLASTx score. We calculated the wGRR between each pair of these genomes, using this as an intergenomic distance metric (1 if each protein has an identical hit in the other genome, and 0 if none of the proteins are similar). We also counted the occurrences of the motif in each genome. Since the motif occurred only in one strand in Rembedalsseter, we calculated the net difference in number of motif occurrences between the two strands of each phage genome (Fig. 8B).

In Fig. 8B, Rembedalsseter clearly clusters with 60 related phages associated with a significantly higher net motif occurrence (9.0 within this cluster vs 0.7 outside this cluster). This cluster includes both Rembedalsseter and Pseudomonas phage tf. Statistically, this motif is also expected to occur by chance 0.7 times in a 45kbp genome, demonstrating that the motif is not enriched in phage genomes outside this cluster. This conserved motif occurrence is not simply a result of high genetic similarity; Rembedalsseter and tf share only minor nucleotide similarity (8% BLASTN coverage), but presumably reflects a conserved function of the nicks. Although it has previously been noted that similar nicked phages share motifs^62^, this is the first systematic investigation into the distribution of nick-associated motifs in sequencing databases, revealing the extent of the motif conservation.

We also investigated how the motif was distributed within these 61 genomes. To enable comparisons between the diverse phages, we reoriented all phage genomes to start from the well-conserved *terL* gene, although this is a different start site than the physical end of the linear genome found in Rembedalsseter (correct orientation shown in Fig. 3D). Fig. 8C shows the nicks are not evenly spread but concentrated towards the right side of the *terL*-oriented genomes. In Pseudomonas phage tf, this part of the DNA molecule has been shown to be the first to attach to phage capsids, prompting the proposal that the nicks may play a role during packaging^62^. Here, we show that this pattern of nick- associated motifs is conserved throughout this group of phages. We also found that only 4/542 motifs were AACT[AG]TGAC, while the remaining were TACT[AG]TGAC, with only three genomes containing AACT[AG]TGAC motifs. However, Rembedalsseter has nicks associated with all four possible versions of the motif.

It has also been suggested that nicks may play a role in phage gene expression^63^. To investigate this, we show the location of motifs relative to the coding sequences predicted by Phanotate (Fig. 8D). While 64 of the 542 motifs occur outside of coding sequences, the rest (88%) occur at least partly within coding sequences. This is nearly identical to the coding density of the host bacteria (89%), indicating coding sequences are not enriched in the motif. Although the motifs often occur at the beginning of coding sequences, they are also found throughout their length. They also occur mostly, though not exclusively, on the coding strand. This distribution throughout coding sequences and strands perhaps argues against a role in gene expression, although this data is merely descriptive and cannot exclude such a role.

We have known about nicked phage genomes for 50 years, yet their function remains enigmatic. Although a role in DNA packaging is plausible, it seems odd that nick-less mutants are viable if they aid such a crucial step in phage reproduction. A role for nicks in gene expression also seems implausible given their distribution both throughout and outside of coding sequences. Perhaps they instead play a role in some non-essential processes, such as homologous recombination (where nicked DNA is an intermediate step) or protection against defense systems (where nicked DNA may protect against sequence recognition). However, all these hypotheses remain untested, and further studies are needed to explain the function of nicked phage genomes.

## 5. Conclusion

In this study, we have isolated eight phages against *Pseudomonas* and *Erwinia* strains from the wheat phyllosphere, contributing to a growing collection of phage isolates targeting wheat phyllosphere bacteria. Based on phylogenetic analysis we propose the creation of four novel phage genera, with two of the phages having no genus-level hits even in metagenomic databases. Contrasting this diversity, other phages appear to be remarkably conserved; the jumbo phage genus *Mimasvirus* appears to be widespread in plant microbiomes, with sequence reads from a near-identical phage to Erwinia phage Kaldavass found in a Spanish spinach microbiome.

Our results also underline the extent to which phages modify their DNA, finding methylation motifs in two jumbo phages and an unknown hypermodification in two *Winklervirus.* We also find a nick-associated motif from Pseudomonas phage Rembedalsseter to be conserved across a group of 61 phages, showing for the first time how ONT sequencing can be used to investigate phage genome nicks. However, the function of these nicks remains unknown. More broadly, these results illustrate how much there is to learn from the isolation and characterization of new phage isolates.

## Acknowledgements

The authors thank Natalie Allcock for her assistance with TEM imaging.

## Funding

This research was supported and funded by the Novo Nordisk Foundation (grant number NNF19SA0059348).

## CRediT authorship contribution statement

**PED**: Investigation, Writing, Visualization, Software, Data Curation, Conceptualization. **MSP**: Investigation, Writing - Review and Editing. **LMF**: Investigation, Software, Visualization, Writing – Review and Editing. **ABC**: Investigation, Writing – Review and Editing. **JMR**: Supervision, Writing – Review and Editing. **LR:** Investigation, Conceptualization, Supervision, Project administration, Writing – Review and Editing. **LHH:** Conceptualization, Supervision, Project administration, Writing – Review and Editing

## Declaration of competing interest

The authors declare that they have no known competing financial interests or personal relationships that could have appeared to influence the work reported in this paper.

## Data availability

The datasets presented in this study can be found in online repositories. The names of the repository/repositories and accession number(s) can be found in the article.

## Notes

### Competing Interest Statement

The authors have declared no competing interest.

